# TDP-43 proteinopathy impairs neuronal mRNP granule mediated postsynaptic local translation and mRNA metabolism

**DOI:** 10.1101/589416

**Authors:** Chia-En Wong, Kuen-Jer Tsai

## Abstract

Local protein synthesis and mRNA metabolism mediated by mRNP granules in the dendrites and the postsynaptic compartments is essential for synaptic remodelling and plasticity in the neuronal cells. Misregulation in these processes caused by TDP-43 proteinopathy lead to neurodegenerative diseases such frontotemporal lobar degeneration (FTLD) and amyotrophic lateral sclerosis (ALS). Using biochemical analysis and imaging techniques including super-resolution microscopy, we provide evidences for the first time of the postsynaptic localization of TDP-43 in the mammalian synapses; and we show TDP-43 as a component of neuronal mRNP granules. With activity stimulation and different molecular approaches, we further demonstrate activity-dependent mRNP granule dynamics involving disassembly of mRNP granules, release of mRNAs, and activation of local protein translation as long as impairments in models of TDP-43 proteinopathy. This study elucidates the interplay between TDP-43 and neuronal mRNP granules in normal physiology and TDP-43 proteinopathy in regulation of local protein translation and mRNA metabolism in the postsynaptic compartment.

## Introduction

Neurodegenerative diseases are a heterogeneous group of disorders characterized by the progressive functional decline of the central nervous system with progressive loss of the structure and the function of neurons in the brain and/or the spinal cord[1][2]. In neurodegenerative diseases including Alzheimer’s disease (AD), frontotemporal lobar degeneration (FTLD), amyotrophic lateral sclerosis ALS and Huntington’s disease (HD), one of the common pathological features is the appearance of abnormal protein aggregates[3][4][5][6][7][8][9]. Previous studies have suggested that the pathogenic process of the abnormal protein aggregates involves abnormal protein folding and can result in loss-of-function of multiple nuclear and/or cytosolic proteins[10]. On the other hand, besides the functional loss caused by abnormal protein aggregation, previously published studies have proven that the formation of insoluble protein inclusions also exerts gain-of-toxicity effects that lead to impaired cellular function and neuronal death in the central nervous system[11][12][13]. Taken together, the processes of abnormal protein folding and aggregation formation contribute both to loss of normal physiological protein function and gain of cellular toxicity, which ultimately lead to disease phenotypes.

RNA-binding proteins(RPBs) are involved extensively in various forms of neurodegenerative diseases[14][15]. RPBs are key components of messenger ribonucleoprotein particles(mRNPs), in which RPBs associate and interact with mRNAs to form protein-mRNA complexes that participated in the regulation of sophiscated post-transcriptional gene regulation including pre-mRNA splicing, RNA editing, mRNA transport and local translation[16]. Recent studies have shown that mRNPs can be further assembled into mRNP granules, which are higher-ordered protein-RNA complexes with the cargo mRNAs associated with non-translating ribosomes. These translationally silenced mRNP granules include neuronal RNA transport granules, P bodies and stress granules[17][18]. Evidences have suggested that the cargo mRNAs in these mRNP granules are translationally paused before completion of the translational initiation and they can transport along axons and dendrites to presynaptic and postsynaptic terminals[19]. Upon appropriate activation signals including synaptic activity, the translationally paused mRNP granules can be further reactivated resulting in local translation in axons, dendrites and synapses[20][21][22]. Among different RBPs, TAR DNA-binding protein 43(TDP-43) is a ubiquitously expressed multifunctional DNA/RNA binding protein that is primarily nuclear but shuttles between the cytoplasm and nucleus[23].TDP-43 was identified as the major component of pathological cytoplasmic inclusions, which are the pathological hallmark of FTLD and ALS[24][25]. Previous studies have shown that both dominant mutations and overexpression of TDP-43 are sufficient to cause FTLD and/or ALS phenotypes, highlighting the role of TDP-43 in disease pathogenesis[26][27][28]. TDP-43 has thousands of mRNA targets, many of which play important role in CNS development and synaptic homeostasis[23], [29]. Despite its primary nuclear localization, cytosolic TDP-43 isolated from both human and rodent brain cytosolic fractions have been shown to bind to the 3’ UTR of numerous target mRNAs[28]. Further studies have also suggested the important role of TDP-43 in mRNA transport and stability. For example, TDP-43 is recognized to be involved in microtubule-dependent bidirectional axonal transport of mRNAs[30]. Somatodendritic TDP-43 was also shown to be regulate in somatodendritic RNA metabolism[31]. Together, these have suggested a potential role of TDP-43 in neuronal plasticity by regulation of axodendritic local translation and impairment of cytosolic TDP-43 function might be of importance of the pathogenesis of diseases featuring TDP-43 proteinopathy such as FTLD and ALS, yet direct evidence is lacking.

Here, we elucidated the interplay between TDP-43 and neuronal mRNP granule in TDP-43 proteinopathy.Here, we provided evidence of the synaptic localization of cytosolic TDP-43 and for the first time directly visualized the postsynaptic localization of TDP-43 in the mammalian synapses by far field super-resolution microscopy. Next, we showed TDP-43 as a component of neuronal mRNP granule and the neuronal activity-dependent granule dynamics in normal physiological condition as well as in TDP-43 proteinopathy. Our work also provided insight into the involvement of neuronal mNRP granule in pathological TDP-43 inclusions in cellular, animal models and FTLD-TDP patients.

## Materials and methods

### Animal models

All the experiments were under review and permitted by the Institute of Animal Use and Care Committee at National Cheng Kung University (NCKU), Taiwan. Mice were all bred at NCKU laboratory animal center. The water and food were provided *ad labitum*. The FTLD-TDP transgenic mouse model carried full-length mouse TDP-43 cDNA under the transcription control of an 8.5-kb CaMKII promoter and overexpressed TDP-43 in the forebrain. Genotyping by PCR was used to identify whether the mice was transgene-positive. At 6 month old, cytosolic TDP-43 aggregation accompanied with nuclear TDP-43 depletion appears in the brain of FTLD-TDP transgenic mice, which mimic the hallmark of TDP-43 proteinopathy in brains of human FTLD-TDP patients[26]. For protein extraction, mice were sacrificed by rapid cervical dislocation and the region of cortex was isolated. For immunofluorescent staining, mice were anesthetized by IP injection of chloral hydrate with the dose of 400mg/kg prior to transcardial perfusion.

### Sub-cellucar fractionation

Subfractionation of mouse brain tissue was performed as described previously with some modifications[32]. Briefly, grossly dissected forebrain tissue is homogenized in buffer (0.32 M sucrose, 20 mM HEPES, pH 7.4, with protease inhibitors) and centrifuged at 1000 × g for 10 min to pellet the membrane fragments and nuclei fraction. The supernatant is collected as the cytosolic fraction, which is then further centrifuged at 17,000 × g for 15 min to obtain the pellet containing crude synaptosome fraction, which are synaptosomes contaminated with mitochondria and microsomes. This crude synaptosome fraction is then further purified using a discontinuous sucrose density gradient consisting of a 1.2 M sucrose layer on the bottom and a 0.8 M sucrose layer on the top. The crude synaptosome suspension is layered on top of the 0.8 M sucrose layer and centrifuged at 54,000 × g for 90 min. The purified synaptosomal fraction is obtained from the interface of 0.8 M sucrose and 1.2 M sucrose.

### Protein extraction

Forebrains containing cortex were resuspended in ice-cold RIPA lysis buffer (50 mM Tris-HCl, pH 8.0, 150 mM NaCl, 1% NP40, 0.5% sodium deoxycholate, 0.1% SDS) containing protease inhibitor cocktail tablet (Roche) and the mixture of phosphatase inhibitor cocktail 2/3 (Sigma Aldrich). Homogenates were centrifuged and supernatants were collected as brain and cell lysates.

### Western blot analysis

Proteins (15-30μg) after quantification by the Bradford protein assay (Bio-rad) were resuspended in loading buffer and subjected to polyacrylamide gel for electrophoresis and transferred to nitrocellulose membrane (Whatman). After blocking, membranes were incubated overnight at 4°C with primary antibodies. Then, membranes were incubated secondary antibodies and developed using Western Lightning^®^ Plus-ECL (PerkinElmer).

### Primary neuronal culture and drug treatment

Before culture, 20-mm glass coverslips were coated with 1 mg/mL of poly-D-lysine at room temperature for 20 minutes; the coated coverslips were then placed into the bottom of 12-well culture plates. Next, mouse pups at postnatal day 0 were sacrificed and their cerebral cortices were excised under a dissection microscope. Cortical tissues were triturated for disaggregation and plated on poly-D-lysine-coated glass coverslips in Neurobasal medium containing B27 serum-free supplement and antibiotics (50 U/mL penicillin and 50 μg/mL streptomycin). Cultures were incubated at 37°C in an atmosphere of 95% air, 5% CO2, and 90% relative humidity. Half of the growth medium was replaced every 2 days. The primary cultured neurons were maintained for 15 to 19 days prior to immunostaining. For repetitive stimulation by KCl [33], the neurons (15 days in vitro) were exposed to repetitive stimuli by KCl. Each stimulus consisted of a treatment with 90 mmol/L KCl for 3 min and a spaced recovery for 10 min. The neuronal cells harvested or fixed for further experiments 30 min after the last stimulation.

### Immunofluorescence staining

For immunohistochemistry of the mouse brains, mice were anesthetized by chloral hydrate and transcardial perfusion was performed with 4% paraformaldehyde (PFA) in PBS. The brain was isolated and then embedded in paraffin blocks. Paraffin section was performed and brains were sectioned at 10-μm-thick for immunofluorescent staining. The sections were deparaffinized in xylene and covered with citrate buffer to perform antigen retrieval. Then permeabilized with 0.5% Triton X-100(Sigma-Aldrich), blocked with 5% bovine serum albumin for 1 hour at room temperature. After blocking, sections were incubated overnight at 4°C with primary antibodies, and then incubated with secondary antibodies for 1 hour. The nuclei were counterstained with DAPI (Sigma-Aldrich). The coverslips were mounted with fluorescent mounting medium.

For immunocytochemistry, cells were washed with PBS and fixed with 4% PFA for 15 min. The cells were then permeabilized with 0.1% Triton X-100 and blocked with 5% bovine serum albumin for 1 hour at room temperature. After blocking, sections were incubated overnight at 4°C with primary antibodies, and then incubated with secondary antibodies for 1 hour. The nuclei were counterstained with DAPI. Images were taken using upright fluorescence microscope (BX51; Olympus) and laser Scanning Confocal Microscope (C1-Si; Nikon).

### dSTORM Microscopy Imaging buffer preparation

We used the imaging buffer composition suggested in a previously reported protocol[34]. Buffer A was composed of 10 mM TRIS (pH 8.0) and 50 mM NaCl, while buffer B was composed of 50 mM TRIS (pH 8.0), 10 mM NaCl, and 10% glucose. GLOX solution (1 mL) was prepared by vortex mixing a solution of 56 mg glucose oxidase, 200 μL catalase (17 mg/mL), and 800 μL of buffer A. MEA solution (1 M, 1 mL) was prepared using 77 mg of MEA and 1 mL of 0.25 N HCl. In each chamber of the 20-mm glass coverslips, 500 μL of imaging buffer was prepared by mixing 5 μL of GLOX solution, 50 μL of MEA solution, and 445 μL of buffer B on ice. After the chamber was filled in, it was covered immediately to avoid replenishing of the dissolved oxygen.

### dSTORM imaging

Alexa Fluor® 647 and Alexa Fluor® 568 secondary antibodies were used in dSTORM imaging. The microscope was constructed around an Olympus IX-83 automated inverted microscope. For illumination, an objective-type, total internal reflection fluorescence, oil-immersion objective (APON 60×; NA 1.49 total internal reflection fluorescence; Olympus) was used. A multiline laser source (405, 488, 561, and 640 nm; Andor Technology) was used for excitation and activation. Single-molecular localization signals were separated using appropriate filters (Andor Technology) and detected using electron-multiplying charge-coupled device camera (iXon Ultra 897; Andor Technology). Before dynamic image movie acquisition, conventional fluorescence images were acquired to determine the region of interest. Specifically, 10,000 frame image series were recorded with an exposure time of 50 ms (20 frames per second). The acquired image series were then analyzed using MetaMorph® Super-Resolution System (Molecular Devices) to generate reconstructed dSTORM images.

### Co-Immunoprecipitation and RNA-Immunoprecipitation

Cells/forebrain tissue were lysed in RIPA lysis buffer (50 mM Tris-HCl, pH 8.0, 150 mM NaCl, 1% NP40, 0.5% sodium deoxycholate, 0.1% SDS) containing protease inhibitor cocktail tablet (Roche) and quantified by the Bradford protein assay (Bio-rad). Proteins (500 μg/1mg) were diluted to 0.5 mg/ml with lysis buffer.

Co-immunoprecipitation assay was performed using the Catch and Release v2.0 Reversible Immunoprecipitation System (Millipore) according to the manufacturer’s instructions with either 4 μg of Anti-TDP-43 antibody (Proteintech) or 4 μg of Anti-FMRP (Cell Signaling Technology). For Western blot analysis, precipitated proteins were washed and subsequently eluted in denaturing buffer and heated at 94°C for 3 min. For RNA-immunoprecipitation analysis, the precipitated proteins were washed and subsequently eluted in non-denaturing buffer, and the RNAs in the immunoprecipitates were purified using Quick-RNA™ Microprep Kit (ZYMO RESEARCH) according to the manufacturer’s instructions.

### Quantitative RT-PCR

For quantitative RT-PCR analysis, the first strand cDNA synthesis was done using the Superscript RT (Invitrogen). Real time PCR was performed using the Power SYBR Green PCR Master Mix (Applied Biosystems) and specific primers for 25–33 cycles of 30 s at 94°C/30 s at 56°C/1 min at72°C. The primers used were: for Map1b, 5’-TGGGACACAAACCTGATTGA-3’ and 5’- TGAAAATCTCTATGAAGTTCT-3’; for GluR1, 5’-CTAGGCTGCCTGAACCTTTG-3’ and 5’-GGGAAGATTGAATGGAAGCA-3’; for CamKII, 5’-AAAGTGCGCAAACAGGAAAT-3’ and 5’-AGGTGGATGTGAGGGTTCAG-3’.

### Ribopuromycilation assay

Ribopuromycilation assay was performed as described previously with some modifications[35][36]. Briefly, primary neuronal cells on coverslips were incubated with Neurobasal medium containing B27 serum-free supplement with 1 mg/ml puromycin and 100 μg/ml cycloheximide for 5 min at 37°C. Afterwards, cells were washed with Neurobasal medium containing B27 serum-free supplement and fixed afterwards for 15 min with 4% PFA in PBS. For immunofluorescence detection, cells were then permeabilized with 0.1% Triton X-100 and blocked with 5% bovine serum albumin for 1 hour at room temperature. After blocking, sections were incubated overnight at 4°C with anti-puromycin antibodies(Millipore), and then incubated with secondary antibodies for 1 hour.

### Urea-soluble fraction preparation

For analysis of urea-soluble proteins, forebrain tissues were dissected and extracted with a serial buffers with increasing reducing power as previously described[37]. Briefly, forebrains were extracted sequentially at 5 mL/g (volume/weight) with low-salt (LS) buffer (10 mM Tris, pH 7.5, 5 mM EDTA, 1 mM DTT, 10% sucrose containing protease inhibitors), high-salt Triton X-100 (TX) buffer [LS buffer + 1% Triton X-100 + 0.5 M NaCl], myelin flotation buffer [TX buffer containing 30% sucrose], and sarkosyl (SARK) buffer [LS + 1% N-lauroyl-sarcosine + 0.5 M NaCl]. The SARK-insoluble fractions were further extracted in 0.25 mL/g urea buffer (7M urea, 2M thiourea, 4% 3-[(3-Cholamidopropyl)dimethylammonio]-1-propanesulfonate, 30 mM Tris, pH 8.5). The urea-soluble proteins were then subjected to Western blotting.

### HEK 293T cell culture and TDP-43 overexpression

Human embryonic kidney (HEK293) cells were maintained in Dulbecco’s modified eagle medium (Gibco) containing 10% fetal bovine serum (HyClone), 1mM sodium pyruvate and 1% penicillin/streptomycin solution (Gibco). HEK293 cells were transfected with GFP-TDP-43 by PolyJet™ DNA *in vitro* transfection reagent (Signagen Laboratories). Cells were seeded into a 6-well plate at a density of 5 × 10^5^ cells/well and incubated overnight. Briefly, 2μg of DNA was diluted to 100 μl with serum-free DMEM with high-glucose (4500 mg/L) (Invitrogen). In another tube, 6 μl of PolyJet reagent was diluted with 94 μl of serum-free DMEM with high-glucose. Diluted PolyJet reagent was added immediately into the diluted DNA solution and gently mixed by pipetting. The mixture was incubated for 15 minutes at room temperature to facilitate the formation of transfection complex. After incubation, DNA/PolyJet complex was added to 6-well plate containing HEK293 cells. After 24 hours, 10 μM of MG-132 (Sigma-Aldrich) was treated for 0, 4, 8 and 12 hours.

### Statistical analysis

All data are reported as the mean ± SEM. The statistical significance of differences between means was evaluated by Student’s *t* test. Differences were considered statistically significant at *p* < 0.05, as indicated by the asterisks.

## Results

### TDP-43 in the Mouse CNS

First, we analyzed TDP-43 distribution in the mouse CNS using Immunofluorescence. We detected TDP-43 in brain areas including the cortical layers and the hippocampus **(Figure S1, a and b).** The sharp-line signals in the hippocampal area indicatea a predominantly somatic localization of TDP-43. In the magnified images, co-staining of TDP-43, neuron marker NeuN and nuclear marker DAPI revealed a predominantly nuclear localization of TDP-43, which was in agreement with previous studies[38][39]**(Figure S1, c).**

### Subcellular Localization of TDP-43

Next, we studied the subcellular localization of TDP-43. Subcellular fractions from forebrain of adult mice were performed. Lamin A and PSD-95 were used as internal control of nuclear and purified synaptosome fraction, respectively. The tissue fractionation showed that TDP-43 was detected in all subfractions including nuclear, cytosolic and purified synaptosome fractions. The result suggested that besides its primary nuclear localization, TDP-43 also showed cytosolic localization and potentially enriched synaptic localization **(Figure 1, a)**.

**Figure 1.**
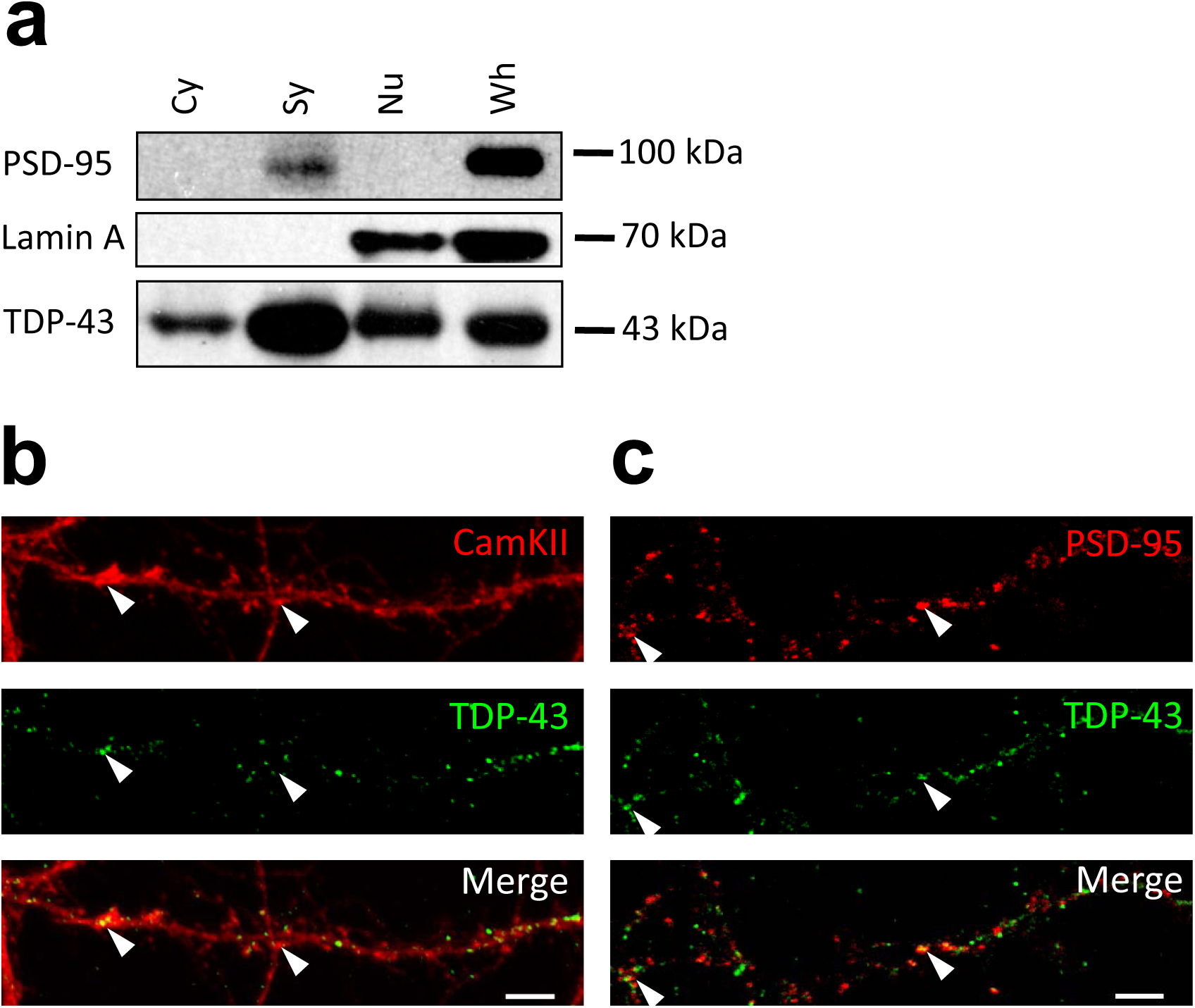
Subcellular Localization of TDP-43. (**a**)Subcellular fractionation from forebrain of adult mice. (**b**,**c**) Confocal microscopy for immunocytochemistry stained TDP-43, CamKII and PSD-95 in primary mouse cortical neuronal cells. Bar: 5µm Arrowheads indicate colocalized TDP-43/CamKII and TDP-43/PSD-95 puncta.

To further confirm the presence of axodendritic and synaptic TDP-43 molecules, we utilized confocal fluorescence microscope to identify the subcellular localization of TDP-43 in 19 DIV primary mouse cortical neurons. We observed a predominant nuclear localization of TDP-43 with cytosolic TDP-43 puncta also present**(Figure S2)**. High magnification images showed that cytosolic TDP-43 puncta distributed along neurites and colocalized with mushroom-like structure of dendritic spines labeled by CamKII**(Figure 1, b)**. Co-staining with synaptic marker PSD-95 also showed colocalization of TDP-43 and PSD-95 puncta in neurites **(Figure 1, c)**. Together, these results suggested that cytosolic TDP-43 molecules have a synaptic subcellular localization in neurons.

### Super-Resolution Microscopy Reveals Postsynaptic Localization of TDP-43 in Dendritic Spines

Hindered by diffraction limit, conventional fluorescence microscopy failed to resolve the exact localization of TDP-43 within cortical synapses. In order to investigate the precise synaptic localization of TDP-43, we utilized super-resolution imaging technique direct stochastic optical reconstruction microscopy (dSTORM). TDP-43 and reference synaptic proteins including presynaptic and postsynaptic makers Bassoon and PSD-95 were immunolabeled in 19 DIV cultured cortical neurons, which contain morphologically mature synapses revealed by presynaptic and postsynaptic makers**(Figure S3)**. Wield field confocal immunofluorescence images showed punca structure of TDP-43, Bassoon and PSD-95 distributed along neuronal processes. However, colocalization of TDP-43 and Bassoon as well as TDP-43 and PSD-95 were both observed in wield field image **(Figure 2, a and c, left panel)**, thus whether TDP-43 localized in presynaptic, postsynaptic or both areas, localization intensity profiles of the protein distributions along the presumed transsynaptic axis unable to determine the exact synaptic localization of TDP-43. Next, we analyzed the reconstructed dSTORM images**(Figure 2, a and c, right panel)**, which showed near-by colocalization of neighboring TDP-43 and Bassoon as well as TDP-43 and PSD-95 in the neuronal processes**(Figure 2, b and d)**. In order to further determine were investigated to measure the transsynaptic axial distance between TDP-43, Bassoon and PSD-95. Here, center of mass of each cluster was determined by fitting to the localization intensity profiles with a simple Gaussian function and the distances between COMs were measured along the transsynaptic axis of the synapses.

**Figure 2.**
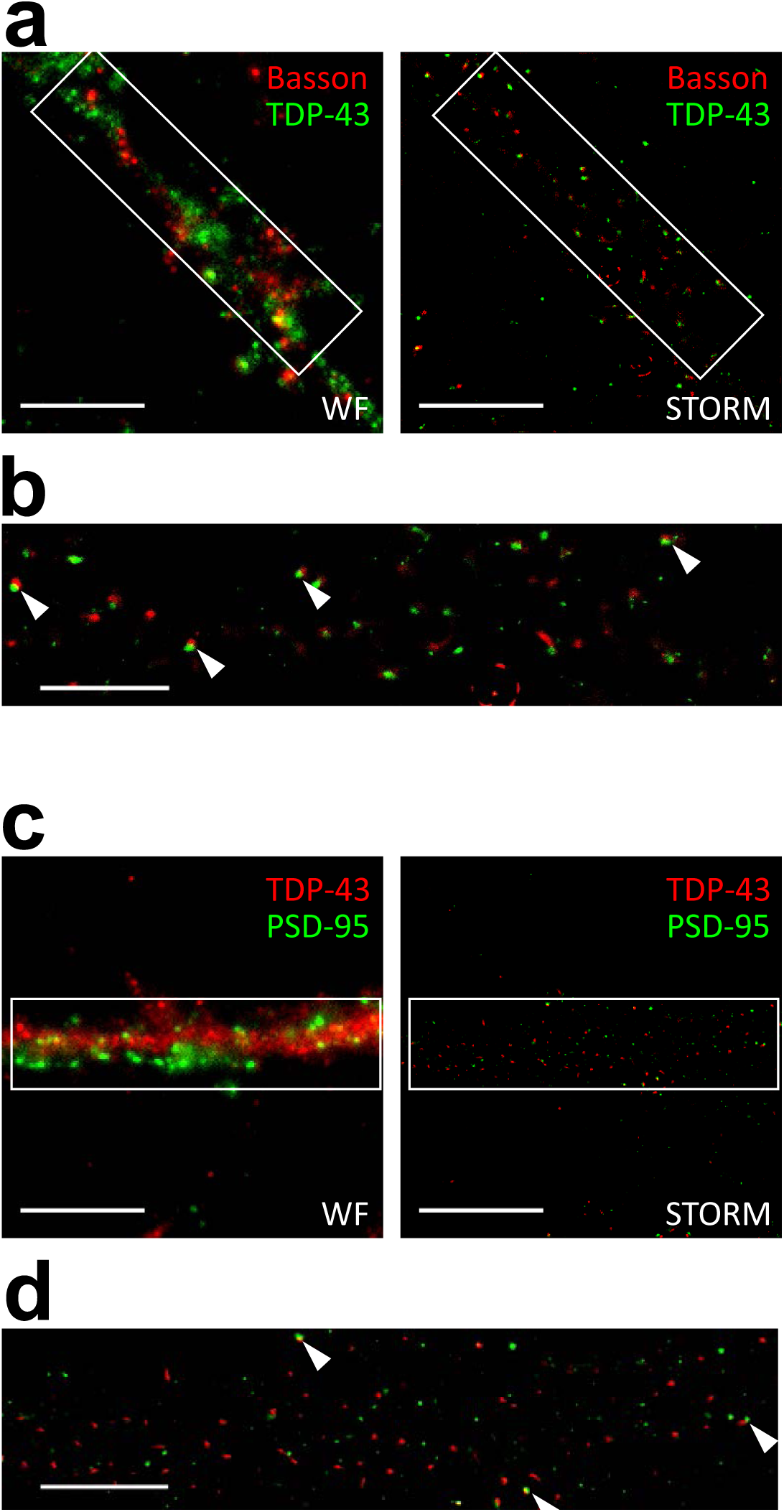
dSTORM images revealed near-by colocalization of synaptic TDP-43 and pre/postsynaptic markers Bassoon and PSD-95. (**a**)Wild-field and dSTORM images of presynaptic marker Bassoon and TDP-43. (**b**)Magnified dSTORM image of Bassoon and TDP-43. Arrowheads indicate near-by colocalization of Bassoon and TDP-43 colo(**c**)Wild-field and dSTORM images of TDP-43 and postsynaptic marker PSD-95. (**d**)Magnified dSTORM image of PSD-95 and TDP-43. Arrowheads indicate near-by colocalization of PSD-95 and TDP-43. Bars in **a**,**c**: 10µm; Bars in **b**,**d**: 5µm

The magnification in **Figure 3** displays typical synapses of each combination of protein cluster pairs of Bassoon and PSD-95, Bassoon and TDP-43, PSD-95 and TDP-43. Position of COMs were calculated from intensity profiles along the transsynaptic axis and the distances of the respective protein pairs were measured **(Figure S2)** As previously described, the following mean distance values arise from imaging a three dimensional structure in two dimensions, as a result, only 40% of all synapses with the highest distance were taken into account to select for synapses whose synaptic cleft is tilted out of and perpendicular to the imaging plan[40]. More than 190 measurements were analyzed and the distances of all analyzed synapses were grouped in a histogram in 20 nm steps to display the distribution of all measurements**(Figure 3, right panels)**.

**Figure 3.**
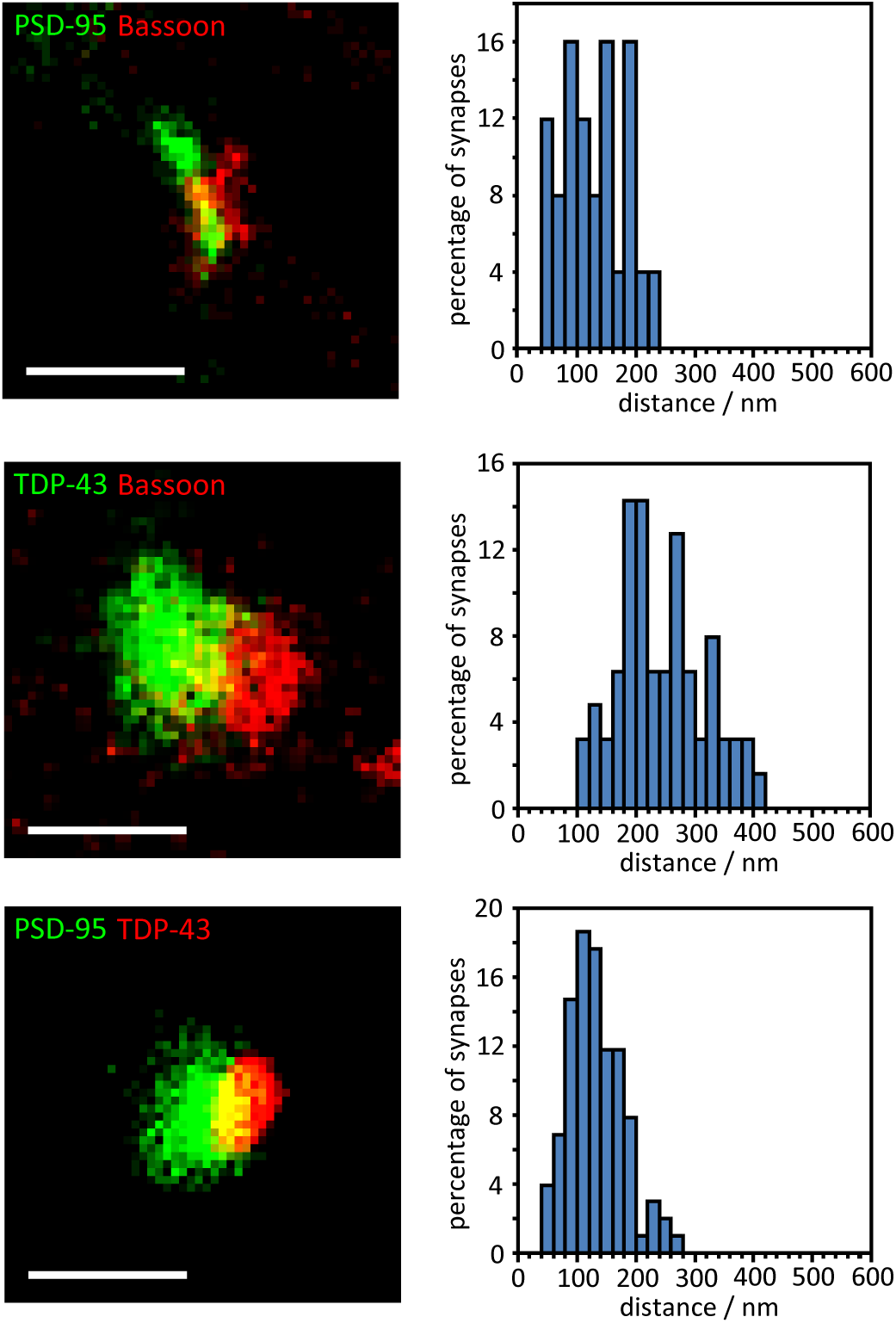
distance analysis revealed postsynaptic localization of synaptic TDP-43. Representative dSTORM images of synaptic pairs of PSD-95, Bassoon and TDP-43(left panels); Histograms showing the distributions of measured distance between PSD-95 and Bassoon, TDP-43 and Bassoon, PSD-95 and TDP-43, respectively.(right panels). Bars: 400nm

The analysis showed a mean distance of 245.1 ± 10.1nm (mean ± SEM, n=63) between TDP-43 and Bassoon, while TDP-43 versus PSD-95 showed a distance of 134.1 ± 5.0nm (mean ± SEM, n=102). The values showed closer proximity of synaptic TDP-43 to the postsynaptic marker PSD-95 compared to the presynaptic marker Bassoon. In addition, the distance between presynaptic marker Bassoon and postsynaptic marker PSD-95 was also analyzed as control measurement, with a mean distance of 121.6 ± 9.5nm (mean ± SEM, n=25), which is in great agreement with previously published data measured in both super-resolution microscopy and electron microscopy.

The distances between TDP-43 and presynaptic/postsynaptic markers lead to two possible position of TDP-43: 1. in the synaptic cleft and closer to the postsynaptic membrane or 2. in the dendritic spine and distal to the postsynaptic density. Importantly, the Bassoon-TDP-43 distance was close to the sum of Bassoon-PSD-95 distance and PSD-95-TDP-43 distance, suggesting a postsynaptic localization of TDP-43 in dendritic spines. Taken together, by combining super-resolved dSTORM microscopy and distance analysis, these results revealed for the first time the postsynaptic localization of synaptic TDP-43 in the dendritic spines.

### TDP-43 as a Component of Neuronal RNA Granules *in vitro* and *in vivo*

Cytosolic TDP-43 binds to numerous target mRNAs in the 3’ UTR region[23]. Previous studies have shown involvement of cytosolic TDP-43 in mRNA transport and stability[30][41]. Some studies have also proposed that TDP- 43 might be involved in neuronal mRNP granules[42][43]. We wonder whether cytosolic TDP-43 interacts with neuronal mRNP granule component as well as markers and whether cytosolic TDP-43 is involved in the regulation of neuronal mRNP granule function and RNA metabolism. To address this, TDP-43 was analyzed versus STAU2 and FMRP. STAU2 and FMRP were both RNA binding proteins and were identified key components and markers of neuronal mRNP granules[44]. Here, 19 DIV primary cortical neurons were cultured and TDP-43, STAU2 and FMRP were immunolabeled. Both STAU2 and FMRP showed puncta distribution in the cytosolic and along the neurites with colocalization of TDP-43 and STAU2 as well as TDP-43 and FMRP observed **(Figure 4, a)**. Next, to further confirm true colocalization of TDP-43 and neuronal granule markers, we utilized super-resolution microscopy. Reconstruction dSTORM images revealed true colocalization of TDP-43 and STAU2 puncta as well as TDP-43 and FMRP puncta **(Figure 4, b)**.

**Figure 4.**
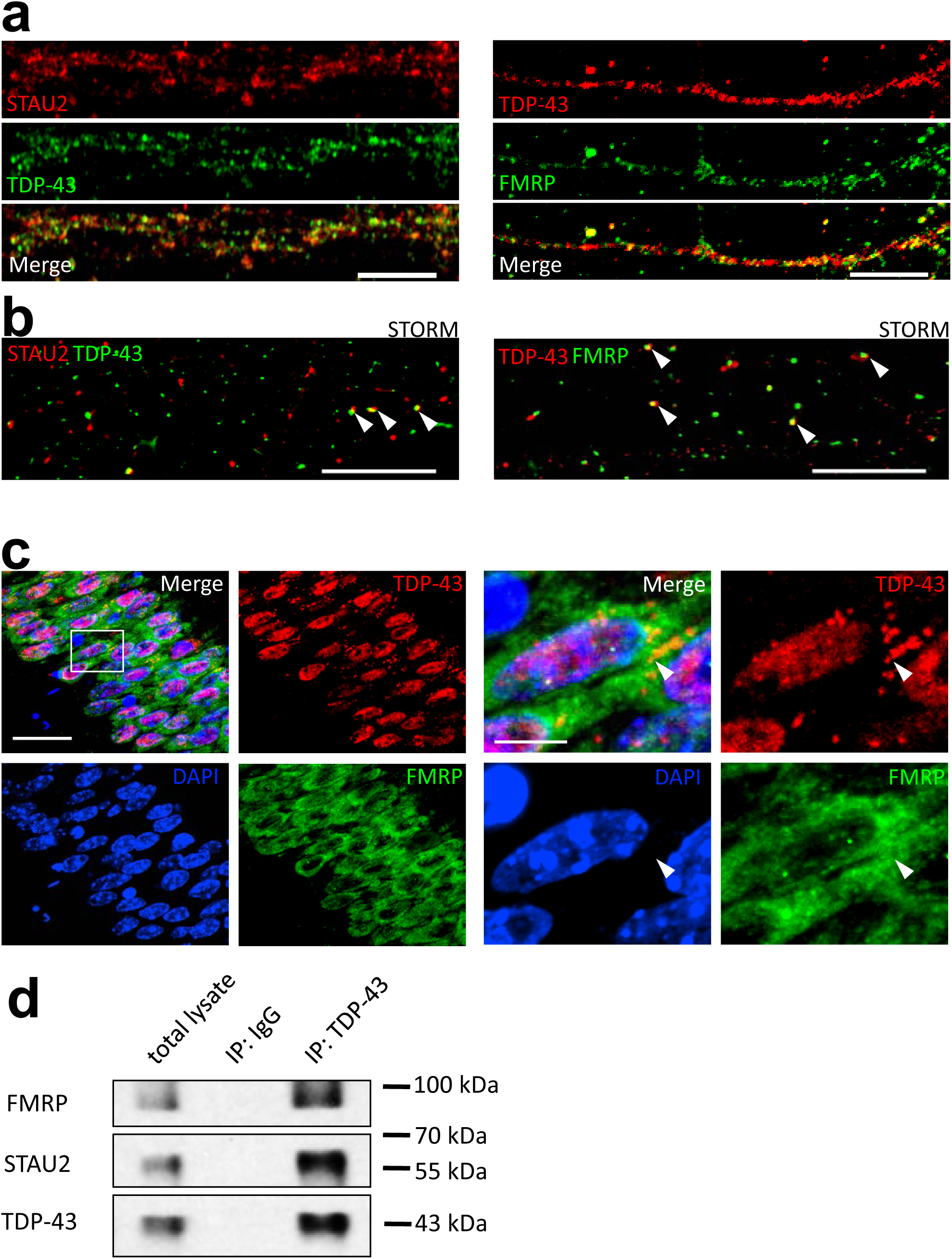
TDP-43 is a Component of Neuronal RNA Granule. (**a**)Confocal microscopy for immunocytochemistry stained TDP-43, STAU2 and FMRP in primary mouse cortical neuronal cells. Bar: 10µm (**b**) dSTORM images of TDP-43, STAU2 and FMRP in primary mouse cortical neuronal cells. Arrowheads indicate colocalized TDP-43/STAU2 and TDP-43/FRMP puncta. Bar: 5µm (**c**)Immunofluorescence staining of TDP-43 and FMRP in adult WT mice hippocampal CA1 area. Arrowheads indicate cytosolic TDP-43 puncta. Bar: 20µm: 5µm (**d**)Co-immunoprecipation for TDP-43 and immuno-blotting for FMRP and STAU2.

To investigate i*n vito* TDP-43, adult mice frozen sections were prepared as previously described method and TDP-43, STAU2 and FMRP were immunolabeled with incubation of primary and secondary antibodies. Confocal microscopy showed a predominant nuclear localization of TDP-43 observed in hippocampal CA1 area**(Figure 4, c, left panels)**. Additionally, cytosolic TDP-43 puncta were also present and colocalization of TDP-43 and FMRP was observed in magnified image**(Figure 4, c, right panels)**.

Besides fluorescent images, biochemical analysis was also performed. We analyzed co-immunoprecipation from whole brain lysates of adult mice. Briefly, whole brain lysates were mixed with resins, anti-TDP-43 antibody and antibody affinity ligand. After incubation for 1 hour at room temperature, the elution was analyzed by western blot. The result showed that there was an interaction between TDP-43, STAU2 and FMRP in adult mice brain**(Figure 4, d)**. Together, these results suggested that cytosolic TDP-43 interacts with neuronal mRNP granule markers and TDP-43 is a component of neuronal mRNP granule.

### Activity-dependent Disassembly of the TDP-43 Containing Neuronal RNA Granule

mRNP granules are dynamic structures and are involved in local translations. Previous studies had shown that different mRNP granules including neuronal granules, processing bodies(PBs) and stress granules(SGs) interact with each other and exchange components[44][45]. More importantly, in the cytoplasm, these translationally silenced mRNP granules can undergo further reactivation for local translation to occur[46]. The reactivation process involves release of paused mRNA-ribosome complex from inhibitory RNA binding proteins that results in initiation of local translation.upon certain signals including neuronal activity[47][48].

Since the binding of TDP-43 to mRNAs has been identified to exert translational inhibition, we hypothesized that upon reactivation by neuronal activity, TDP-43 containing mRNP granules are disassembled and TDP-43 are unbound from the mRNP complexes so that local translation can be completed.

To address this, we performed stimulation on 15 DIV primary mouse cortical neurons by repeated depolarization. Briefly, the neurons were treated by 90 mmol/L KCl three times, each time with 3 minute duration and 10 minute spaced recovery between each treatment. The neurons were fixed 30 minutes after the last treatment. We analyzed the distribution of TDP-43 and FMRP on mouse cortical neurons by immunofluorescence with or without KCl stimulation. In comparison to the unstimulated neurons, neurons treated with KCl showed less colocalized puncta of TDP-43 and neuronal mRNP granule marker FMRP along the neurites**(Figure 5, a, left panel)**. Furthermore, the confocal images of the neurites of unstimulated and stimulated neurons were analyzed by Pearson’s correlation coefficient and object based colocalization analysis. A one third reduction of colocalization of TDP-43 with neuronal mRNP granule marker FMRP was observed in both Pearson’s correlation coefficient and object based colocalization analysis**(Figure 5, b)**. These results suggested that the interaction between TDP-43 and neuronal mRNP granules were decreased upon stimulation, which supported our hypothesis that TDP-43 containing mRNP granules are disassembled and TDP-43 are unbound from the mRNP complexes upon neuronal activity.

**Figure 5.**
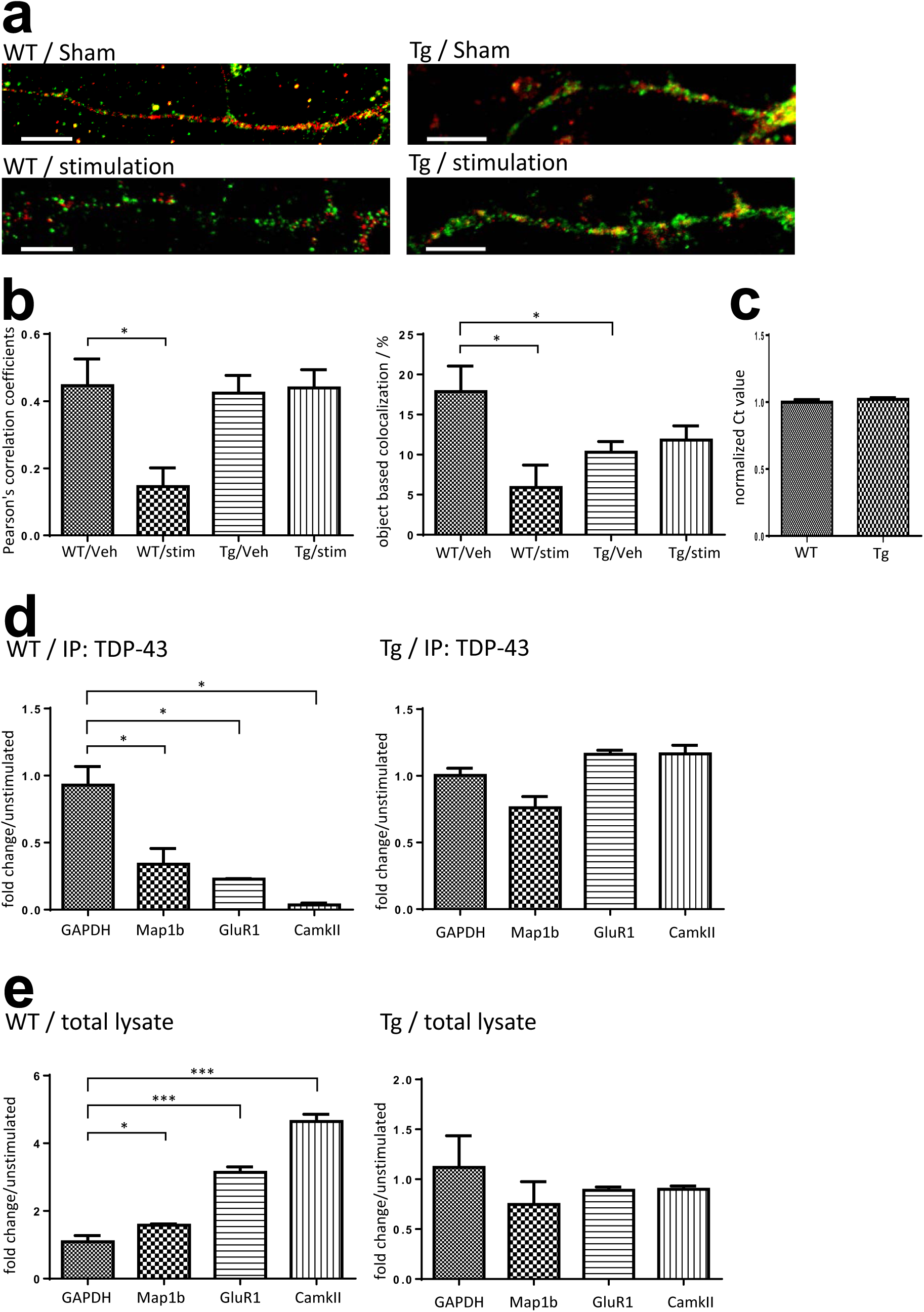
Activity-dependent Dynamics of TDP-43 Containing Neuronal RNA Granule and its impairment in TDP-43 proteinopathy. (**a**)Confocal microscopy for immunocytochemistry stained TDP-43 and FMRP in WT/Tg primary mouse cortical neurons 30 minutes after KCl stimulation or sham treatment. Bar: 10µm (**b**)Pearson’s correlation coefficient and object based colocalization analysis of TDP-43 and FMRP in stimulated and unstimulated WT/Tg neurons. (**c**)Normalized GADPH RNA level in WT and Tg neurons. (**d**)Fold change of TDP-43 co-immunoprecipated RNA levels in stimulated WT/Tg neurons compared to unstimulated neurons. (**e**)Fold change of free form RNA levels in stimulated WT/Tg neurons compared to unstimulated neurons.

### Impairment of Activity-dependent Granule Disassembly in TDP-43 Proteinopathy

Since that TDP-43 is involved extensively in neuronal mRNP granule metabolism and that TDP-43 proteinopathy is sufficient to cause neurodegeneration and FTLD phenotypes, we wondered whether TDP-43 proteinopathy may lead to impair neuronal mRNP granule function. We used a TDP-43 overexpression transgenic mouse model(TDP-43 Tg) which features FTLD phenotype including impaired learning/memory function and formation of cytosolic TDP-43 inclusions. We had generated a FTLD-TDP mouse model that specifically overexpresses full-length TDP-43 in the forebrain under the control of the Ca2+/calmodulin-dependent kinase II (CamKII) promoter[26]. The FTLD-TDP mice featured impaired performances in the learning/memory tests and formation of cytosolic TDP-43 inclusions.

We first investigated whether the activity-dependent mRNP granule disassembly and TDP-43 unbinding were affected in TDP-43 proteinopathy. Here, 15 DIV primary mouse cortical neurons were prepared from P0 pups of TDP-43 Tg mice and stimulation was performed according to the same protocol of repeated KCl depolarization as that performed on WT neurons. The unstimulated and stimulated TDP-43 Tg neurons were then analyzed by immunofluorescence and confocal microscopy to investigate the distribution of TDP-43 and FMRP**(Figure 5, a, right panel)**. The result showed that in comparison to the unstimulated Tg neurons, there were no obvious difference in colocalization of TDP-43 and FMRP in the KCl treated Tg neurons. Further analysis by Pearson’s correlation coefficient and object based colocalization also confirmed that there was no difference between the stimulated and unstimulated Tg neurons in colocalization**(Figure 5, b)**. These results suggested that the activity dependent mRNP granule disassembly was impaired in Tg neurons with TDP-43 proteinopathy.

Next, in order to investigate whether the activity dependent mRNP granule disassembly and its impairment in TDP-43 proteinopathy are related to TDP-43-mRNA interactions and RNA metabolism in TDP-43 containing mRNP granules, we investigated the profiles of TDP-43 target mRNAs in the unstimulated and stimulated neurons. Previous studies have identified many mRNA targets of TDP-43, among which some are involved in regulation of synaptic strength and neuronal plasticity[29][49]. For example, Map1b, GluR1 and CamKII are involved in dendritic spine remodelling and regulation of synaptic signaling[50][51][52]. We thus investigated the quantity of TPD-43 bound mRNAs and free form mRNAs. RNA-immunoprecipitation(RNA-IP) were performed from stimulated and unstimulated 15-DIV neuron lysates with anti-TDP-43 antibody and RNA extracts were analyzed by quantitative PCR (qPCR) to detect mRNP granule bound form mRNAs. In addition, free form mRNA extracted from 15-DIV neuron lysates with TRIzol reagent were also analyzed by qPCR. GAPDH mRNA was used as internal control to normalize the RNA levels. There was no difference in baseline GAPDH mRNA level between WT and TDP-43 Tg neurons**(Figure 5, c)**.

The qPCR results from anti-TDP-43 immunoprecipitated WT neurons stimulated by KCl showed decreased levels of TDP-43 bound target mRNAs with 0.34, 0.23 and 0.04 fold changes of Map1b, GluR1 and CamKII mRNA levels normalized to GAPDH compared to the unstimulated WT neurons**(Figure 5, d, left panel)**. Importantly, the decrease of TDP-43 bound mRNAs after stimulation was observed with concurrent increase of free form Map1b, GluR1 and CamKII mRNA levels with 1.58, 3.14 and 4.64 fold changes compare to the unstimulated WT neurons respectively**(Figure 5, e, left panel)**. Together these results supported the idea that the activity dependent mRNP granule disassembly resulted in release of mRNP bound mRNAs, which became free form mRNAs.

However, the activity dependent shift from mRNP bound mRNAs to free form mRNAs was not observed in TDP-43 Proteinopathy. Both the qPCR results of anti-TDP-43 immunoprecipitated mRNAs and free form RNAs from TDP-43 Tg neurons showed no difference between the unstimulated and the stimulated Tg neurons in all Map1b, GluR1 and CamKII mRNA levels**(Figure 5, d and e, right panels)**. This further suggested that the impairment of activity dependent mRNP granule disassembly in TDP-43 proteinopathy resulted in subsequent impairment of mRNA release from the TDP-43 containing mRNP granules.

### Impairment of mRNP Granule Mediated Local Translation in TDP-43 Proteinopathy

Next, in order to determine whether the activity-dependent mRNP granule disassembly and subsequent mRNA release were linked to changes in local translation in the dendrites, we conducted ribopuromycilation assay to measure active translation in stimulated and unstimulated WT/Tg neurons. The experimental procedures of ribopuromycilation assay were described in Materials and methods section.

In the neurons, puromycilated puncta were observed distributing along the neurites.**(Figure 6, a)** In WT neurons, quantification showed that stimulation by KCl significantly increased the number of puromycin labeled puncta, suggesting an increased local translation in the dendrites induced by neuronal activity**(Figure 6, b)**. Compared to WT neurons, TDP-43 Tg neurons showed a lower basal translation. Additionally, in TDP-43 Tg neurons, the activity dependent increase of local translation was not observed and there was no difference in dendritic local translation between KCl stimulated and unstimulated neurons baring TDP-43 proteinopathy**(Figure 6, b)**.

**Figure 6.**
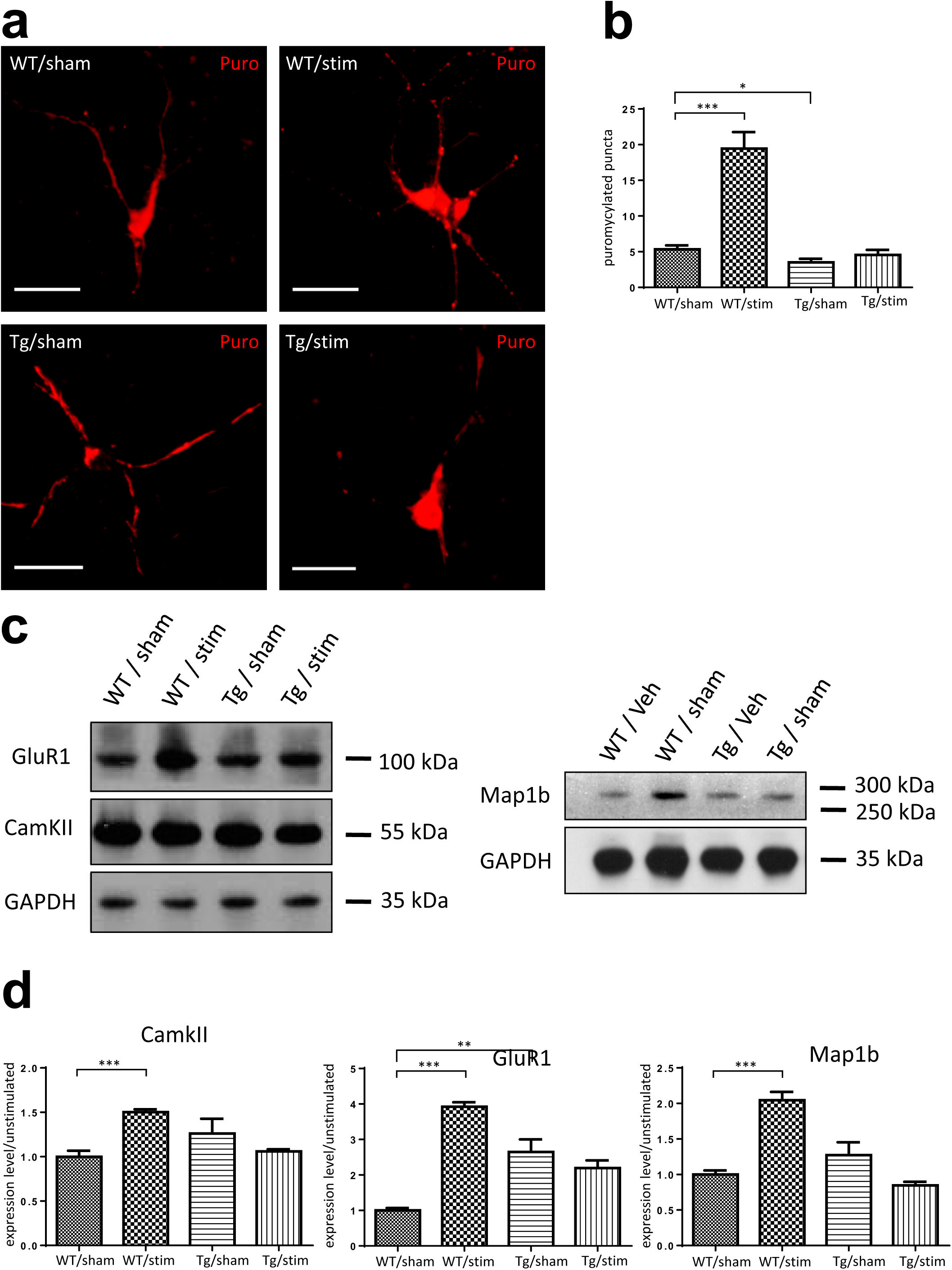
mRNP Granule Mediated Local Translation is impaired in TDP-43 Proteinopathy. (**a**)Immunocytochemistry of ribopuromycilated WT/Tg neurons receiving KCl or sham treatments. Bar: 20µm. (**b**)Quantification of ribopuromycilated puncta per cell in WT/Tg neurons receiving KCl or sham treatments. (**c**)Western-blotting for GluR1, CamKII and Map1b in WT/Tg neurons receiving KCl or sham treatments. (**d**) Quantification of Western-blot protein levels for GluR1, CamKII and Map1b.

Furthermore, we analyzed protein level of the above discussed TDP-43 target proteins Map1b, GluR1 and CamKII in unstimulated and KCl stimulated WT/Tg neurons. Western blots were performed with GAPDH as internal control**(Figure 6, c)**. Quantification of the protein levels was performed using ImageJ software. The result showed that in WT neurons, the KCl stimulated neurons had 2.04 fold, 3.92 fold and 1.50 fold increased relative protein levels of Map1b, GluR1 and CamKII compared to the unstimulated neurons, respectively. In contrast, there was no difference in TDP-43 target protein levels between Tg neurons with and without stimulation**(Figure 6, d)**.

Together the RPM results and the western blot results suggested that the above described granule disassembly and subsequent mRNA release were linked to increased local translation in dendrites and increased levels of TDP-43 target proteins. Importantly, the result further confirmed our hypothesis that the neuronal granule mediated RNA metabolism and local dendritic translation were impaired in model of TDP-43 proteinopathy. In summary, we reported activity-dependent mRNP granule disassembly accompanied by subsequent mRNA release, increase in local translation, increased levels of TDP-43 target proteins and their impairments in TDP-43 proteinopathy.

Additionally, comparing the unstimulated WT and Tg neurons, we also reported a difference in basal protein level of GluR1, indicating an increased amount of glutamate AMPA receptor in TDP-43 Tg neurons. This might imply the potential disruption of glutamate receptors caused by TDP-43 proteinopathy where the functional synaptic deficit in excitatory neurons were compensated by up-regulation of AMPA receptors. Supporting evidences included previous studies showing disruption of glutamate receptors in models of neurodegeneration[53][54].

### Dysregulated RNA Granules as a Component of Pathological TDP-43 Inclusions in FTLD Mouse Model

We elucidated the metabolism of TDP-43 containing mRNP granule, however, the role of neuronal mRNP granule in pathological TDP-43 inclusions remained unclear. To further elucidate the role of neuronal mRNP granule in pathological TDP-43 inclusions, we investigated whether mRNP granule markers were present in TDP-43 pathological inclusions. First, co-immunoprecipation with anti-TDP-43 antibody showed interaction between TDP-43 and neuronal mRNP granule markers FMRP and STAU2**(Figure 4, d)**. Additionally, co-immunoprecipation from 19 DIV WT/Tg neurons performed with anti-FMRP antibody showed increased relative amount of TDP-43 interacting with FMRP in TDP-43 Tg neurons compared to WT neurons**(Figure 7, a)**. Next, to direct visualize the TDP-43 increase, dSTORM images of immunolabled TDP-43 and FMRP were investigated, in which TDP-43 containing neuronal mRNP granules were identified as colocalizing puncta of TDP-43 and FMRP **(Figure 7, b, left panel)**. The quantification revealed increased TDP-43 area and increased TDP-43 total intensity per FMRP/TDP-43 colocalized puncta in Tg neurons compared to WT neurons**(Figure 7, b, right panel)**, which suggested increased relative amount of TDP-43 interacting with FMRP in Tg neurons compared to WT neurons and was in agreement with the result of co-immunoprecipation.

**Figure 7.**
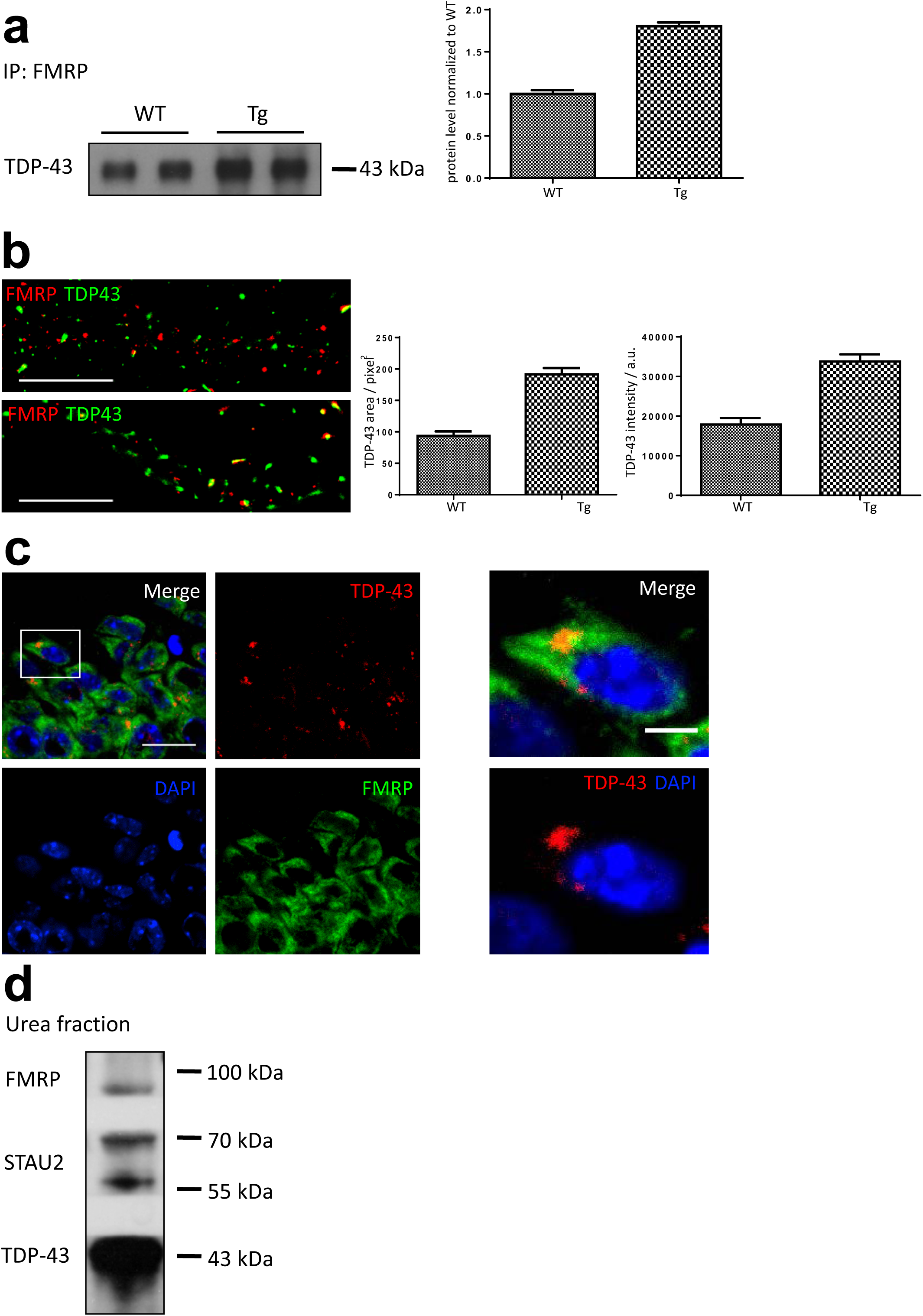
Neuronal mRNP Granule is a Component of pathological TDP-43 inclusion. (**a**)Co-immunoprecipation for FMRP and immuno-blotting for TDP-43(left panel); Quantification of the co-IP TDP-43 level. (**b**)dSTORM images of TDP-43 and FMRP in WT and Tg primary mouse cortical neuronal cells(left panel) Bar: 5µm; Quantification of TDP-43 area and TDP-43 intensity. (**c**)Immunofluorescence staining of TDP-43 and FMRP in adult TDP-43 Tg mice hippocampal CA1 area. Bar: 20µm: 5µm (**d**)Western-blotting for TDP-43, STAU2 and FMRP in urea soluble fractions.

Furthermore, to investigate the role of neuronal mRNP granule in pathological TDP-43 inclusions *in vivo*, frozen sections from 1y FTLD mouse brain featuring TDP-43 proteinopathy were analyzed by immunofluorescence. In contrast to the predominant nuclear localization of TDP-43 in WT mouse brain sections**(Figure S1, c)**, the immunofluorescence images of TDP-43 Tg mice brain showed predominant cytosolic TDP-43 puncta distribution in the cytoplasm of hippocampal CA1 neurons. There were little colocalization between TDP-43 and DAPI, indicating the cytosolic puncta distribution was accompanied by a nuclear clearance of TDP-43 **(Figure 7, c, left panel)**. Additionally, colocalization of TDP-43 and FMRP was also observed in the magnified image **(Figure 7, c, right panel)**.

Next, we utilized the technique of urea fractionation to purify the water-insoluble urea-soluble TDP-43 inclusions from 1y FTLD mouse brain. The urea fractionation was analyzed by western blot, in which neuronal mRNP granule markers STAU2 and FMRP were both co-purified within the water-insoluble urea-soluble TDP-43 fraction indicating that neuronal mRNP granule components were present in pathological TDP-43 inclusions **(Figure 7, d)**.

### Involvement of Neuronal mRNP Granule in the Formation of TDP-43 Cytosolic Inclusions

Since our evidences had suggested that neuronal mRNP granules were involved in pathological TDP-43 inclusions, we further investigated whether mRNP granules were present and involved during the process of formation of the pathological TDP-43 inclusions. Here we used a cellular model to study the formation of TDP-43 inclusions. TDP-43-GFP was transfected by lipofectamine and overexpressed in HEK 293T cell line. After transfection for 24 hours, protease inhibitor MG-132 was added to the cell culture to allow overexpression and formation of cytosolic TDP-43 inclusions. The HEK293-T cells were observed by confocal microscopy at 0, 4, 8 and 12 hours after MG-132 treatment. As control group, there were no TDP-43-GFP signal in HEK 293-T cells with no transfection and vehicle transfection **(Figure S5, left panels)**. On the other hand, the TDP-43-GFP transfected cells showed positive TDP-43 signal at all time points including 0, 4, 8 and 12 hours after MG-132 treatment **(Figure S5, right panels)**. Next, we observed that cytosolic TDP-43 was not present in HEK cell observed immediately after MG-132 treatment and the percentage of cells with cytosolic TDP-43 signal among all transfected cells (defined as cells with TDP-43 positive signal, regardless of the subcellular localization) increased in a time-dependent manner**(Figure 8, b, left panel)**.

**Figure 8.**
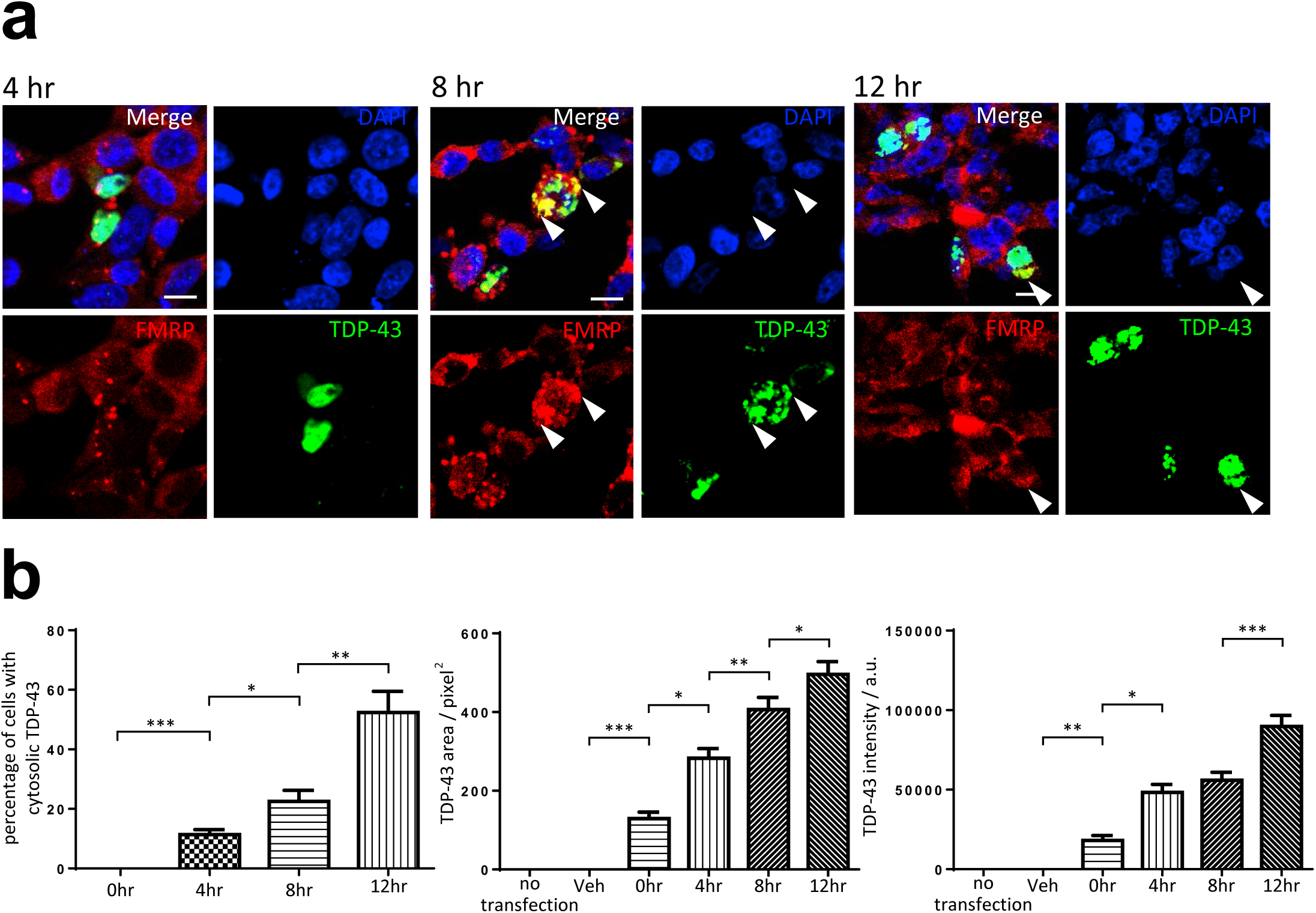
mRNP Granule is Involved in the Formation of TDP-43 Cytosolic Inclusions. (**a**)Immunocytochemistry staining of FMRP in GFP-TDP-43 transfected cells after overexpression for 4, 8 and 12 hours. Arrowheads indicate cytosolic TDP-43 puncta. Bar: 20μm. (**b**) Percentage of cells with cytosolic TDP-43(left panel); Quantification of TDP-43 area and intensity per transfected cell(middle and right panel).

In addition, TDP-43 inclusion bodies presenting as dense cytosolic puncta were observed in HEK 293-T cells after 8 and 12 hours of MG-132 treatment**(Figure 8, a, middle and right panels)**. We further analyzed the average TDP-43 area and total TDP-43 intensity per transfected cell. The analysis showed both increases in TDP-43 area and total TDP-43 intensity per transfected cell in a time-dependent manner in the cells, which revealed the accumulation of cytosolic TDP-43 over time during the process of inclusion body formation**(Figure 8, b, middle and right panels)**. Importantly, we also reported that colocalization between mRNP granule marker FRMP and TDP-43 were observed in all time points when cytosolic TDP-43 were present in 4, 8 and 12 hours after treatment**(Figure 8, a)**, further suggesting the involvement of mRNP granule component during the accumulation of cytosolic TDP-43 over time.

Taken together, these results suggested not only mRNP granules were present in TDP-43 pathological inclusions, but they were also involved during the process of TDP-43 inclusion formation.

### Role of Neuronal mRNP Granule in Pathological TDP-43 Inclusions in the Brain of FTLD-TDP Patients

Our evidence elucidated the role of neuronal mRNP granule in pathological TDP-43 inclusions, where mRNP granules not only were present in TDP-43 pathological inclusions, but also they were involved during the process of TDP-43 inclusion formation. However, these experiments were performed in animal or cellular models. Thus we wonder whether similar evidence could be observed in a more clinically relevant model.

We investigated the cerebral cortical sections obtained from five FTLD-TDP patients featuring TDP-43 proteinopathies. Immunofluorescence revealed FTLD-TDP pathological hallmarks with dense cytosolic TDP-43 aggregations accompanied by clearance of nuclear TDP-43 in the cerebral cortex of all five FTLD-TDP patients**(Figure 9, upper and middle panels)**. Importantly, counterstaining of neuronal mRNP granule marker FMRP showed high-degree colocalization between FMRP signals in the cytosolic TDP-43 aggregates in the cerebral cortex of all five FTLD-TDP patients**(Figure 9, lower panels)**. This result further supported the idea that the involvement of neuronal mRNP granule in TDP-43 proteinopathy and pathological TDP-43 inclusions shown in animal and cellular experiments was relevant and reproducible in true FTLD-TDP patients.

**Figure 9.**
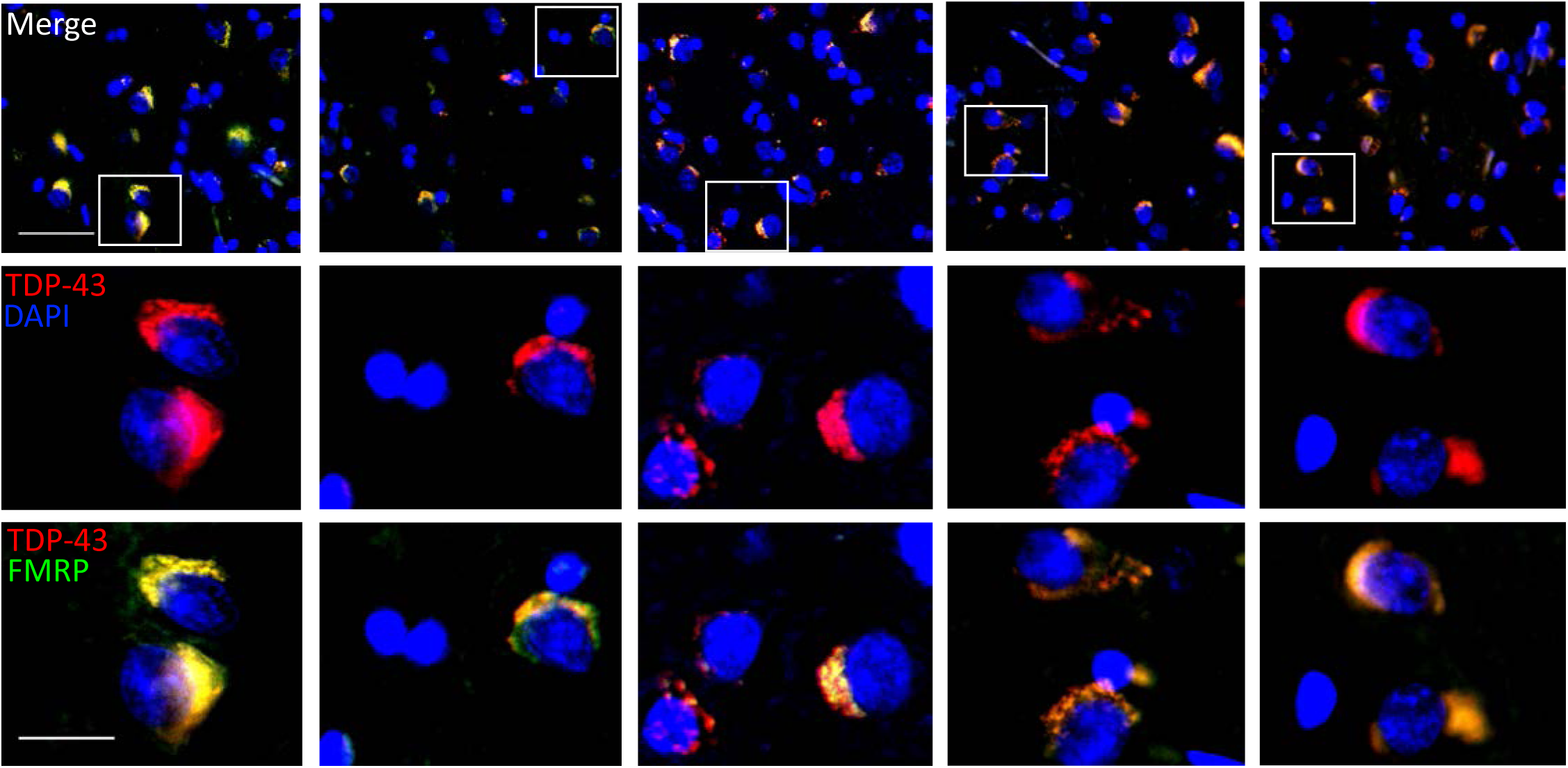
Role of neuronal mRNP granule in TDP-43 proteinopathies in FTLD-TDP patients. Immunofluorescence staining for TDP-43 and FMRP in the frontal cortex of five FTLD patients.

## Discussion

A detailed elucidation of the interplay between TDP-43 and neuronal mRNP granule in TDP-43 proteinopathy might impact neuron biology and would increase our understanding of the molecular pathogenic mechanisms of neurodegenerative disorders and of the metabolism of neuronal RNP granules. Our work provided several important insights into these issues.

First, the biochemical fractionation experiments and immunofluorescence here supported that besides the predominant nuclear localization, cytosolic localizations of TDP-43 in neurons were also present and were compatible with previous reported results regarding the subcellular localization of TDP-43 with a predominant nuclear localization along with cytosolic distributions[55][56]. Further investigation using super-resolved dSTORM microscopy revealed close proximity with near-by colocalization of neighboring TDP-43 and pre/postsynaptic markers. In addition, distance analysis here provided evidence for the first time the postsynaptic localization of synaptic TDP-43 in the dendritic spines. Since TDP-43 is a RNA/DNA binding protein and serves as translational repressor[57], its localization in the dendritic spines might suggest potential role of TDP-43 in the regulation of synaptic local RNA metabolism.

As previous studies had suggested involvement of TDP-43 in neuronal mRNP granule and might exert activity-dependent dynamics[58][30], we reported TDP-43 as neuronal mRNP granule component. Importantly, the observations reported here suggested potential mechanism of TDP-43 in regulation of local synaptic translation. We showed the dynamics of TDP-43 containing neuronal mRNP granules; upon neuronal activity, mRNP granules were disassembled and downstream local translation of TDP-43 mRNA targets was induced. These downstream mRNA targets including Map1b, GluR1 and CamKII all playing critical roles in synapse remodeling[52][59][60], also highlighting the crucial of TDP-43 regulation in synaptic plasticity.

Our work revealed that TDP-43 proteinopathy perturbed the activity-dependent dynamics of TDP-43 mRNP granules and resulted in impaired local translation of plasticity related mRNAs. In addition, the excessive accumulation of TDP-43 in neuronal mRNP granules was also reported here. Since previous studies had shown that intermolecular interactions of TDP-43 via C-terminal low complexity domain resulted in tight bounding potentially causing the formation of pathological inclusions[61][62], it is quite probable that the tight bounding of neuronal mRNP granule component resulted from excessive accumulation of TDP-43 might hinder the disassembly of neuronal mRNP granule and impair subsequent mRNA release and local translation. Evidence here revealed potential mechanism of TDP-43 proteinopathy related impairment of synaptic plasticity, which resulted in impaired learning and memory in FTLD[26][63]

Given the accumulation of TDP-43 in neuronal mRNP granule, it is quite probable that mRNP granules with TDP-43 accumulation might serve as seeds for more TDP-43 to accumulate and eventually to form pathological inclusions. This was supported by our result showing the presence of mRNP granule components both in TDP-43 pathological inclusions and during the process of inclusion formation as well. Importantly, we also confirmed the pathological phenotype in human FTLD-TDP patient brains.

Taken together, these results proposed a model of FTLD pathogenesis, in which the TDP-43 misregulation in the postsynaptic density first caused perturbation of normal neuronal mRNP granule function and lead to impairment of local translation dependent synaptic plasticity and decline of learning memory function. At later stage of disease, the TDP-43 accumulated mRNP granule further served as seeds during the formation of pathological inclusions, which were reported to exert neurotoxicity leading to neuronal death[64][65].

Looking forward, it will be important to further investigate the molecular mechanism of TDP-43 in regulation of activity dependent neuronal mRNP granule dynamics of normal and pathological conditions. Understanding the basis for these processes may provide potential therapeutic strategies to the synaptic dysfunction in early stage FTLD. More importantly, as we have shown, mRNP granule may serves as potential seeds to facilitate formation of pathological inclusions. Further knowledge about the process of pathological inclusion formation may reveal further therapeutic targets to prevent or reverse the formation of TDP-43 pathological inclusions.

The observations reported here are likely to pertain to TDP-43 related neurodegenerative diseases including FTLD and ALS. In summary, our work not only confirmed the cytosolic and dendritic localization of TDP-43 but showed an additional and precise localization of the protein at the postsynaptic density in primary mouse cortical neurons. We revealed the involvement of TDP-43 as a component of neuronal mRNP granule and in the regulation of activity dependent granule dynamics and local dendritic translation. In cellular, animal and human model of TDP-43 proteinopathy, impairment of the neuronal mRNP granule function and the involvement of neuronal mRNP granules in the formation of pathological inclusions were also reported. Our work here provided insights into the pathogenesis and potential therapeutic targets of FTLD.

---

**Figure S1.**
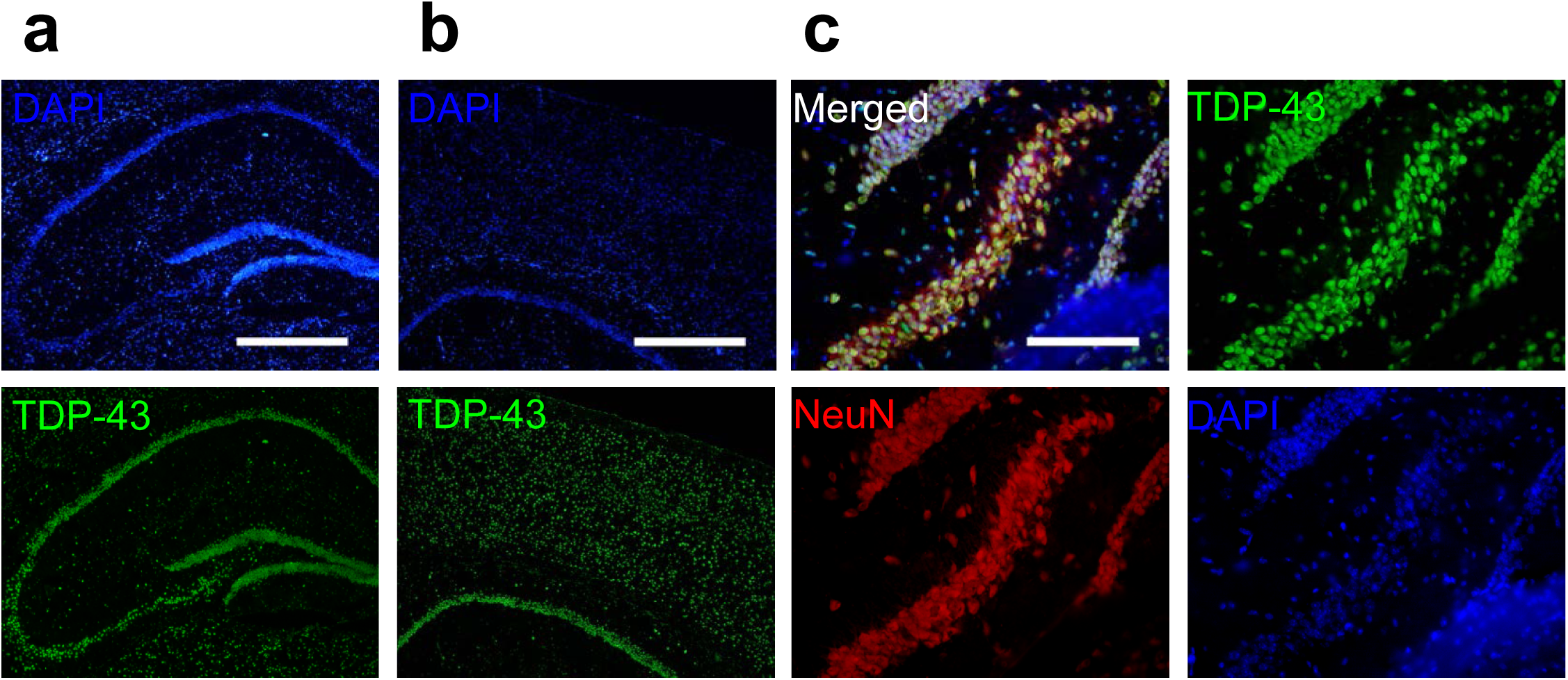

**Figure S2.**
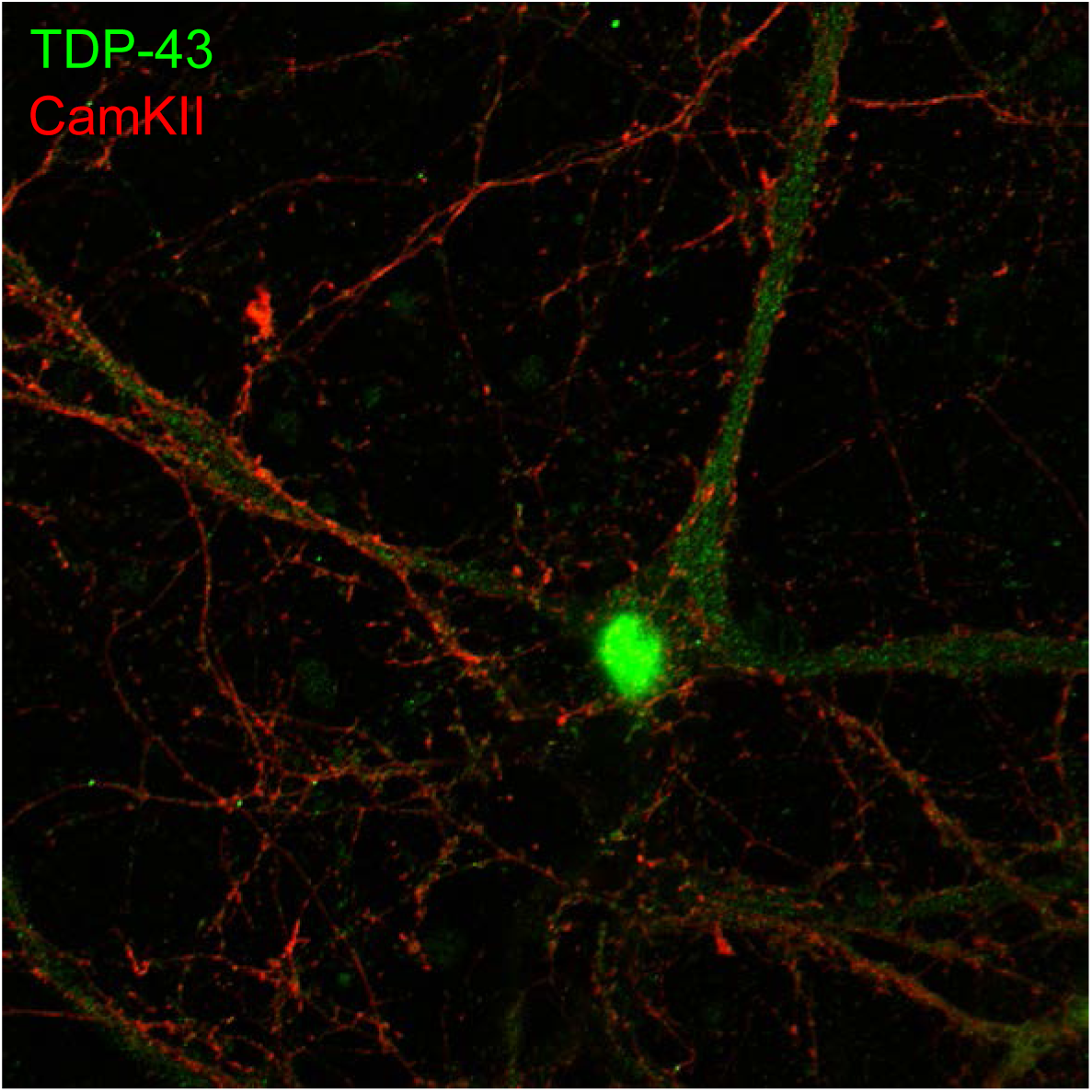

**Figure S3.**
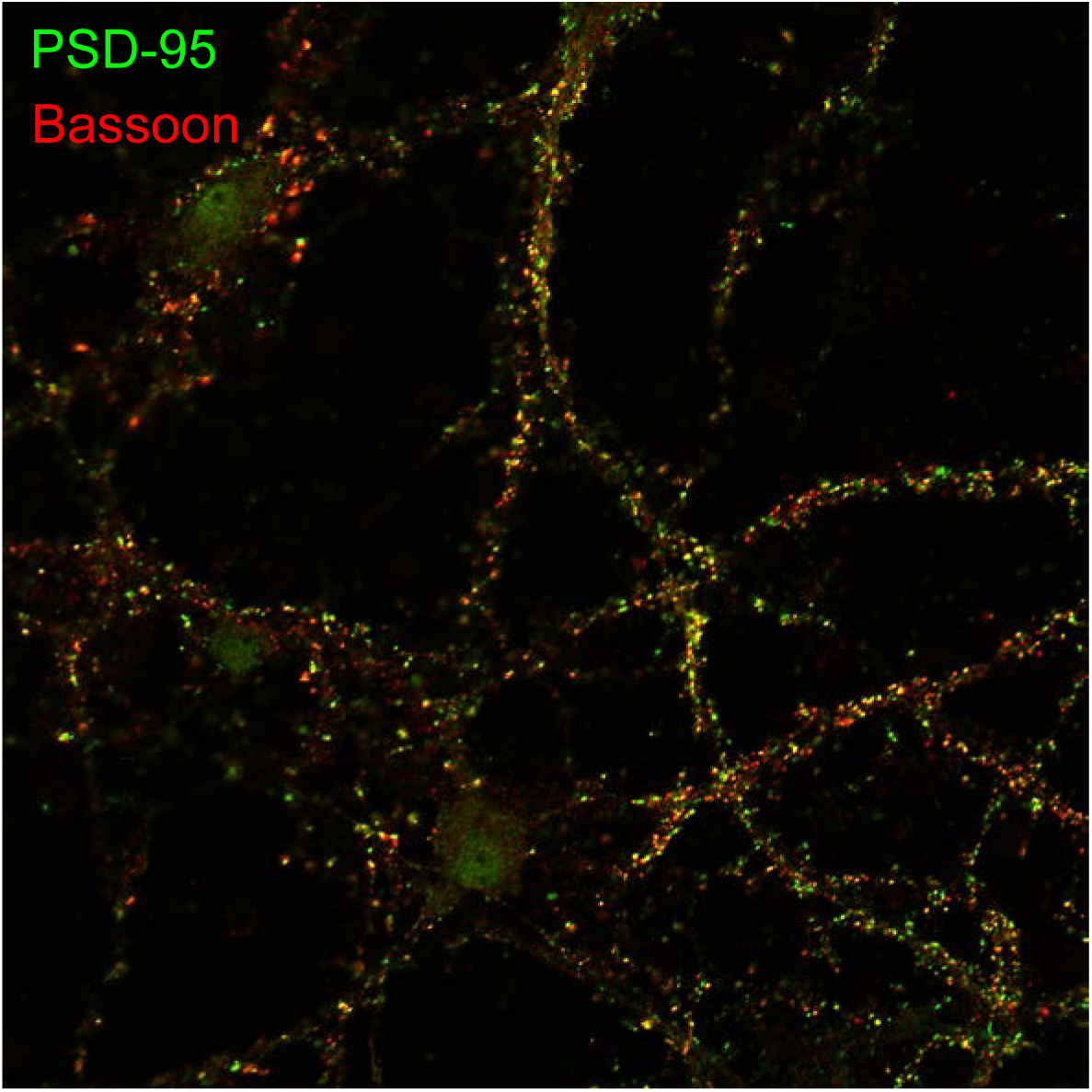

**Figure S4.**
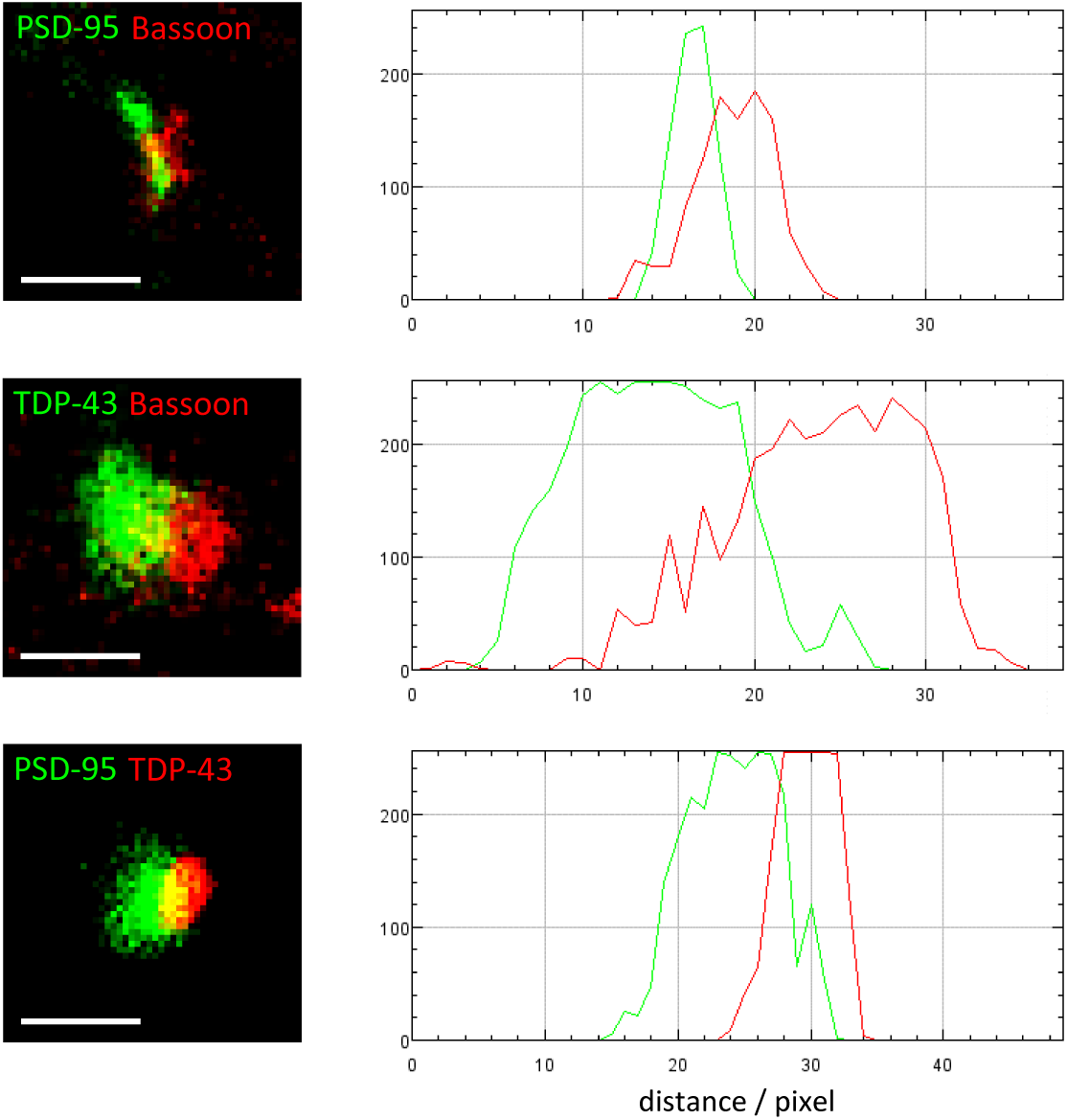

**Figure S5.**
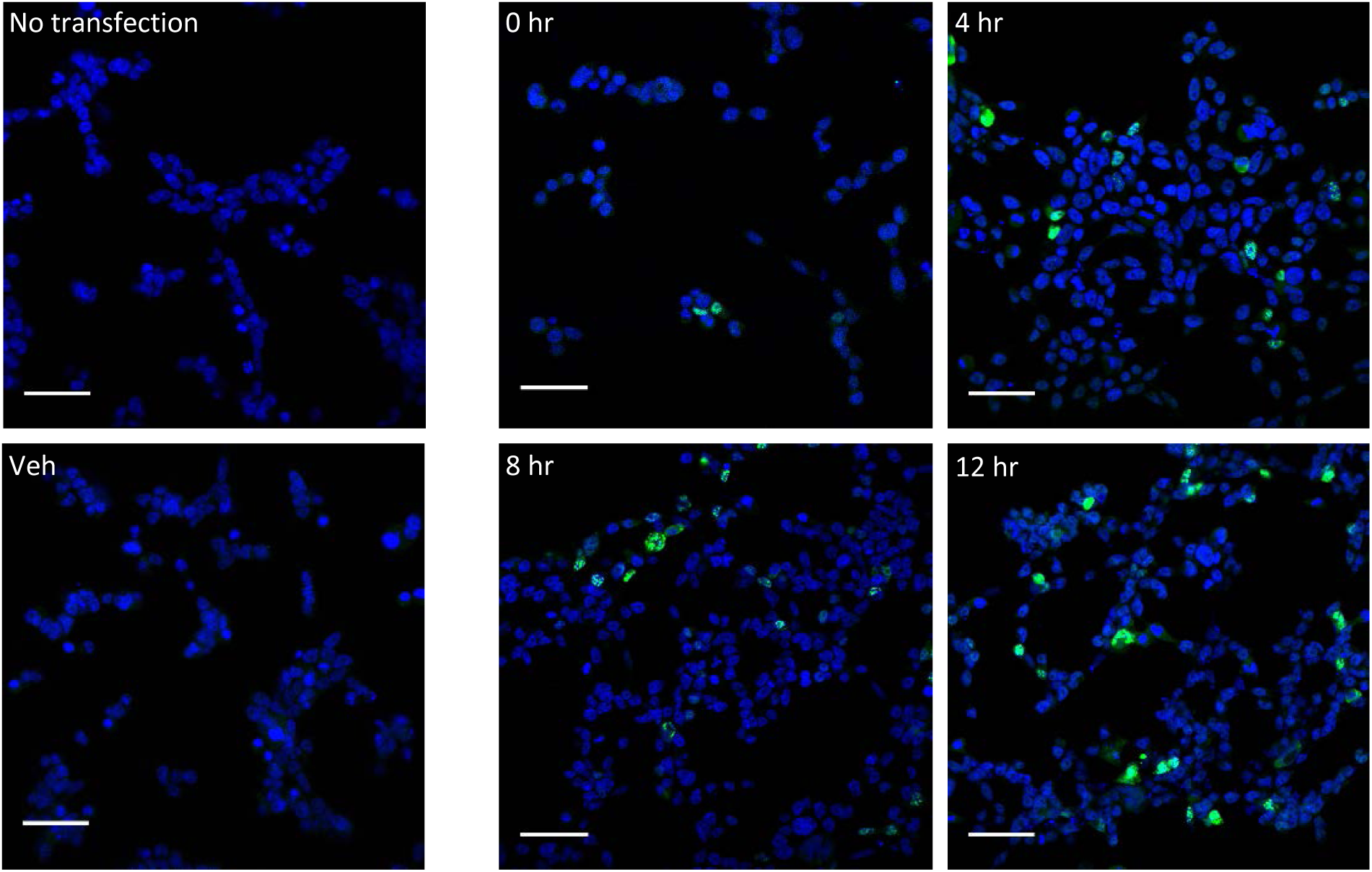

